# Phylogenetic inference from single-cell RNA-seq data

**DOI:** 10.1101/2022.09.27.509725

**Authors:** Xuan Liu, Jason Griffiths, Isaac Bishara, Jiayi Liu, Andrea H. Bild, Jeffrey T. Chang

## Abstract

Tumors are comprised of subpopulations of cancer cells that harbor distinct genetic profiles and phenotypes that evolve over time and during treatment. By reconstructing the course of cancer evolution, we can understand the acquisition of the malignant properties that drive tumor progression. Unfortunately, recovering the evolutionary relationship of individual cancer cells linked to their phenotypes remains a difficult challenge. To address this issue, we have developed PhylinSic, a method that reconstructs the phylogenetic relationships among cells linked to their gene expression profiles from single cell RNA-sequencing (scRNA-Seq) data, and showed that it was robust to the low read depth, drop-out, and noisiness of scRNA-Seq data. This method called nucleotide bases from scRNA-Seq reads using a probabilistic smoothing approach, and then estimated a phylogenetic tree using a Bayesian modeling algorithm. We evaluated PhylinSic and showed that it identified evolutionary relationships resulting from selective events such as drug selection and metastasis and was sensitive enough to identify subclones from genetic drift. Finally, we applied methods of phylogenetic inference and found that breast tumors resistant to chemotherapies harbored two genetic lineages that independently manifested high predicted activity of K-Ras and β-catenin, potentially acquired by distinct mechanisms through convergent evolution. This suggested that therapeutic strategies may need to target multiple lineages to be durable. Taken together, these results demonstrated that PhylinSic provides a framework to model the evolution and link the genotypes and phenotypes of cells within a tumor or cohort of monophyletic tumors using scRNA-Seq.

## Introduction

Since it was first documented by Nowell [1], the cancer cell evolution model where selective pressures produce cancer cell subpopulations with malignant traits has gained an increasing amount of support [2, 3]. Subpopulations of cancer cells, or subclones, could share distinct genotypes as well as phenotypes [4], and may exhibit different malignant properties including proliferation rates, drug resistance, or metastatic capacity. Therefore, having an understanding of the evolution of individual cancer cells in a tumor would help to reveal the traits that are under genetic selection, how they are acquired, and also potential targets to inhibit them.

To uncover the genetic architecture of cancer cells in a tumor, cell barcoding strategies have been developed [5-8]. For samples without barcodes, early methods have been applied to next generation sequencing of the DNA (genome, exome, or select targets) from bulk tumors, where frequencies of subclones were inferred based on the frequencies of mutations seen in the reads [9, 10]. For more precise resolution than could be obtained by bulk sequencing, approaches using single-cell DNA sequencing (scDNA-Seq) have also been explored [11]. However, while these methods could reconstruct the evolutionary histories of cancer cells, they did not reveal the phenotypes of the cancer cells. To study the phenotypes, methods have been developed that could infer linkages between scDNA-Seq and scRNA-Seq (single-cell RNA sequencing) data in samples where profiling were performed in parallel (but on different cells) [12]. Although single-cell multi-omic protocols to sequence DNA and RNA from the same cell have been developed, this strategy has remained technically challenging [13-15].

To find subclones from only scRNA-Seq data, a common approach was to predict copy number variation from gene expression measurements, and then to perform hierarchical clustering on the copy numbers [16, 17]. Although the resultant dendrograms resembled a phylogeny, they did not reflect an underlying model of evolution and thus, interpreting them as speciation events required the assumption that the metric of distance between copy numbers reflected the time needed to acquire the mutations. Without such a model of the molecular clock, or a model of noise in the measurements, it was difficult to interpret the history or groupings of the cells as evolutionary events. As an alternative approach, an early pre-print had demonstrated the feasibility of generating phylogenies using mutation calls from scRNA-Seq [18]. Unfortunately, validation was limited. We found that performance was constrained by the difficulty of determining nucleotide base calls as most commonly used scRNA-Seq technologies suffered from low coverage and high drop-out rates [19].

To address the challenges in inferring phylogenies from scRNA-Seq data, we have developed a method, PhylinSic (<u>Phyl</u>ogeny <u>in Si</u>ngle <u>c</u>ells, bred in a pandemic and pronounced “feeling sick”), that called bases by borrowing information from genetically similar cells when coverage was low, and then applied a Bayesian phylogenetic inference algorithm BEAST2 that could model the imputed base calls [20]. We evaluated PhylinSic on a data set of ER+ breast cancer cells where drug-resistant and -sensitive lineages were fluorescently labeled. We found it robust to noise and able to reconstruct cancer cell phylogenetic relationships and uncover how phylogeny was reflected in cell phenotypes. We applied PhylinSic to investigate drug resistance in breast cancer and multiple myeloma tumors, showing its applicability across biological contexts. In all data sets, we found evidence of genetic evolution across disease progression, as well as evidence for convergent evolution where multiple lineages evolved toward a shared mechanism of resistance.

## Methods

### Generating alignments

We started by generating a BAM file containing the aligned reads for each cell. For the 10X scRNA sequencing platform, we used the standard CellRanger pipeline, which generated a BAM file (*possorted*.*bam*) that contained the alignments for all the cells in each sample. We demultiplexed this file based on the error-corrected, confirmed barcodes for each cell in the CB tag (ignoring the GEM well suffix) in the alignment lines. To speed up the genotype calls, we discarded low coverage cells with less than 10,000 total reads.

### Calling genotypes

We next identified the genomic positions where nucleotide bases differed across cells. To do this, we combined the demultiplexed BAM files into a single pseudobulk sample comprised of monophyletic cells (i.e., from the same person). The variants from the pseudobulk sample were then called using the GATK RNA-Seq pipeline [21, 22]. In short, we added read groups, split reads, recalibrated base quality scores, realigned indels, and applied HaplotypeCaller. We filtered for the variants that were supported by at least 20 reads (across all cells), with at least five variant reads making up at least 5% of the total reads. This resulted in a list of the genomic sites that varied across the cells in the sample, candidates for subclonal mutations.

Next, for each of the variant sites, we extracted the number of reference and alternate allele reads from the cell-specific BAM files, resulting in two parallel matrices. One contained the number of reference reads for each of the sites (rows) and cells (columns). The other contained the number of alternate allele reads.

Using the pair of reference and alternate read count matrices, we modeled the reference and alternate allele read counts as a probabilistic distribution over the alternate (or variant) allele frequency to account for the noise in the scRNA-Seq data. We used a beta-binomial distribution where the number of reference and alternate alleles were observed (i.e., the number of alternate alleles were successes in a Bernoulli trial).

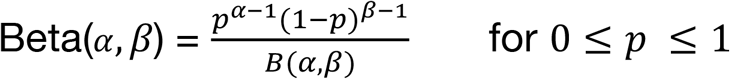

where *p* was the variant allele frequency, *α*-1 was the number of alternate reads and *β*-1 was the number of reference reads.

Based on the distribution of alternate allele frequency, we called the genotypes for each site in every cell. We distinguished among three possible genotypes: homozygous for the reference allele, homozygous for the alternate allele, or heterozygous. While theoretically, a reference genotype should only have yielded reference reads and an alternate genotype should only have yielded alternate reads, in practice, technical noise led to erroneous reads. To account for noise in the data, we introduced a parameter *θ* and obtained the probability that the underlying genotype was homozygous reference by integrating over the probability density function from 0 to *θ* (homozygous reference) or 1-*θ* to 1 for a homozygous alternate genotype, and the remaining probability was allocated to the heterozygous genotype. The value of *θ* depended on the amount of noise in the data (which was difficult to measure), and we used *θ*=0.3. We assigned a genotype by choosing the one with the highest probability.

### Smoothing genotypes

Due to the low read depth frequently seen in scRNA-Seq, most genotype calls were supported by few reads and were therefore uncertain. To improve the reliability of the calls, we adopted a smoothing strategy where we borrowed information from similar cells. We used a kNN approach where we identified cells with similar mutation profiles, and then calculated the average of their probability distributions, weighted by read depth.

To find the neighbors of each cell, we used a mutation similarity score between each cell and all other cells. To calculate this score, we started by estimating the probability distribution over the three genotypes for each cell and site using the beta-binomial distribution, as described above. Then, for each cell, we sampled from the distribution for each site to yield a randomly simulated profile of the genotypes. To score the similarity of the genotype profiles between two cells, we used a Jaccard index (the percent of sites with the same genotype). The final similarity score between a pair of cells was the average Jaccard index over 100 samplings.

Using the pairwise similarity scores of each cell, we chose the *K* nearest neighbors for each cell. We smoothed the probabilities for each genotype by:

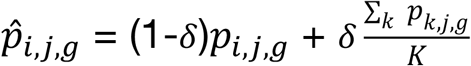

where *p*_*i,j*_ was the probability that cell *i* and site *j* had genotype g (either reference, alternate, or heterozygous). Since *K* was the number of neighbors used for smoothing, it should be less than the size, in cells, of the smallest clade of interest. *δ* was the fraction of probability mass to assign to the neighbors, and higher *δ* resulted in more smoothing. By default, we used *K*=10 and *δ* = *K*/(*K*+1).

We have made the code for genotype calling and smoothing available at: https://github.com/U54Bioinformatics/PhylinSic_Project/blob/main/smoothing.R

### Filtering Sites

Using the reference and alternate allele count matrices, along with the smoothed genotype calls, we filtered for high quality sites. In general, we first removed sites that 1) had low overall coverage (cells with <10 reads), 2) had the same base in more than 90% of cells, 3) were located outside chromosomes 1-22, and 4) were within five nucleotides of another site. Next, we smoothed the genotypes as described above. Finally, we kept the 1000 sites with the highest number of reads. We used more forgiving thresholds (removed sites where cells had less than five reads, and had the same base in more than 80% of cells) for the CAMA-1 data set (see below) because it had relatively poor coverage. For the breast cancer data set, we smoothed genotypes using five neighboring cells due to the low number of cells.

We annotated the genomic metadata (e.g. genomic region, gene affected, impact on protein translation, etc) for all remaining sites using Annovar [23].

### Generating and analyzing phylogenies

We formatted the filtered genotype matrix as a FASTA file for phylogenetic analysis. Each cell was converted into a sequence record containing the bases at each site. At homozygous sites, we used the reference or alternate bases observed in the sequencing data. For heterozygous sites, we chose a base other than the reference or alternate arbitrarily.

To model the phylogenetic relatedness of cells, we used BEAST2 with a relaxed log-normal clock model (RLN), generalized time-reversible site model (GTR) and Yule tree priors. We ran each Markov chain Monte Carlo (MCMC) chain for at least 100 million iterations and added iterations if convergence was not seen in the predicted phylogenies as visualized by a densitree, or by a plateau in the posterior probability of the model. We also monitored the mutation probabilities and rates. We summarized the model as a maximum clade credibility (MCC) tree.

### Phylogenetic measurements

The heritability of cancer traits was measured by the phenotypic resemblance of closely related cells. For each of the 50 Hallmark pathways [24], we used Pagel’s λ measure of phylogenetic signal to quantify the association between the pathway scores from ssGSEA and the phylogenetic structure. This measure of phylogenetic heritability was calculated using the phylosig function in the Phytools R package [25].

We defined the phenotype signal as the ability to distinguish evolutionarily divergent cancer genotypes (known resistant and sensitive lineages) based on their RNA expression profiles. We quantified this by performing a UMAP clustering of the transcriptomic profile of all cells and used the adjusted random index to measure the distinctness of the phenotype of the genotypes using the adjustedRandIndex function implemented in the R package GeometricMorphometricsMix.

### Evolutionary diversity

The evolutionary diversity within a group of cells was measured by their mean pairwise (cophenetic) distance (MPD) across the phylogeny. The difference in evolutionary diversity of multiple groups of cancer cells (e.g. pre- vs post-treatment samples) was measured by the standardized effect size (SES) of the MPD within each group of cells, and was calculated using the ses.mpd function in the R package picante [26]. We identified samples with significantly more or less evolutionary diversity than would be expected by chance by randomly sampling from the phylogeny using a bootstrap permutation test.

### Gene expression analyses

The expression profiles of the single cells were processed using Seurat [27]. The count matrix was normalized using the LogNormalize method and the 3000 most variable genes (selection.method=vst) were identified. Principal component analysis (PCA) of the variable genes provided 50 principal components and UMAP (uniform manifold approximation and projection) further reduced the data into 10 dimensions. Next, the cells were clustered with SNN (shared nearest neighbors) in 10 dimensions with resolution=0.5. Finally, we identified genes with log fold change > 0.05 between resistant and sensitive cells and are expressed in >5% of cells. We removed these differentially expressed genes and verified by visualization in a UMAP plot that this lenient fold change cutoff led to the elimination of the differences in the gene expression profiles of the resistant and sensitive cells.

To characterize the phenotype of the cancer cells, we performed ssGSEA (single-sample gene set enrichment analysis) [28] to obtain signature scores of the Hallmark pathways from the Molecular Signatures Database [24, 29]. The pathway scores were normalized using a zero-inflated negative binomial model to account for the zero inflated (dropout) and over-dispersed read count data [30].

## Results

### Modeling phylogenies from scRNA-Seq data

We developed an algorithm to estimate the evolutionary relationships among monophyletic cells (e.g., those derived from the same person) profiled with scRNA-Seq technologies. The algorithm had three major steps: 1) identification of variant sites, 2) inferring the nucleotide base at the variant sites for each cell, using smoothing to account for scRNA-seq noise, and then 3) reconstructing the phylogeny (**Figure 1A**).

**Figure 1.**
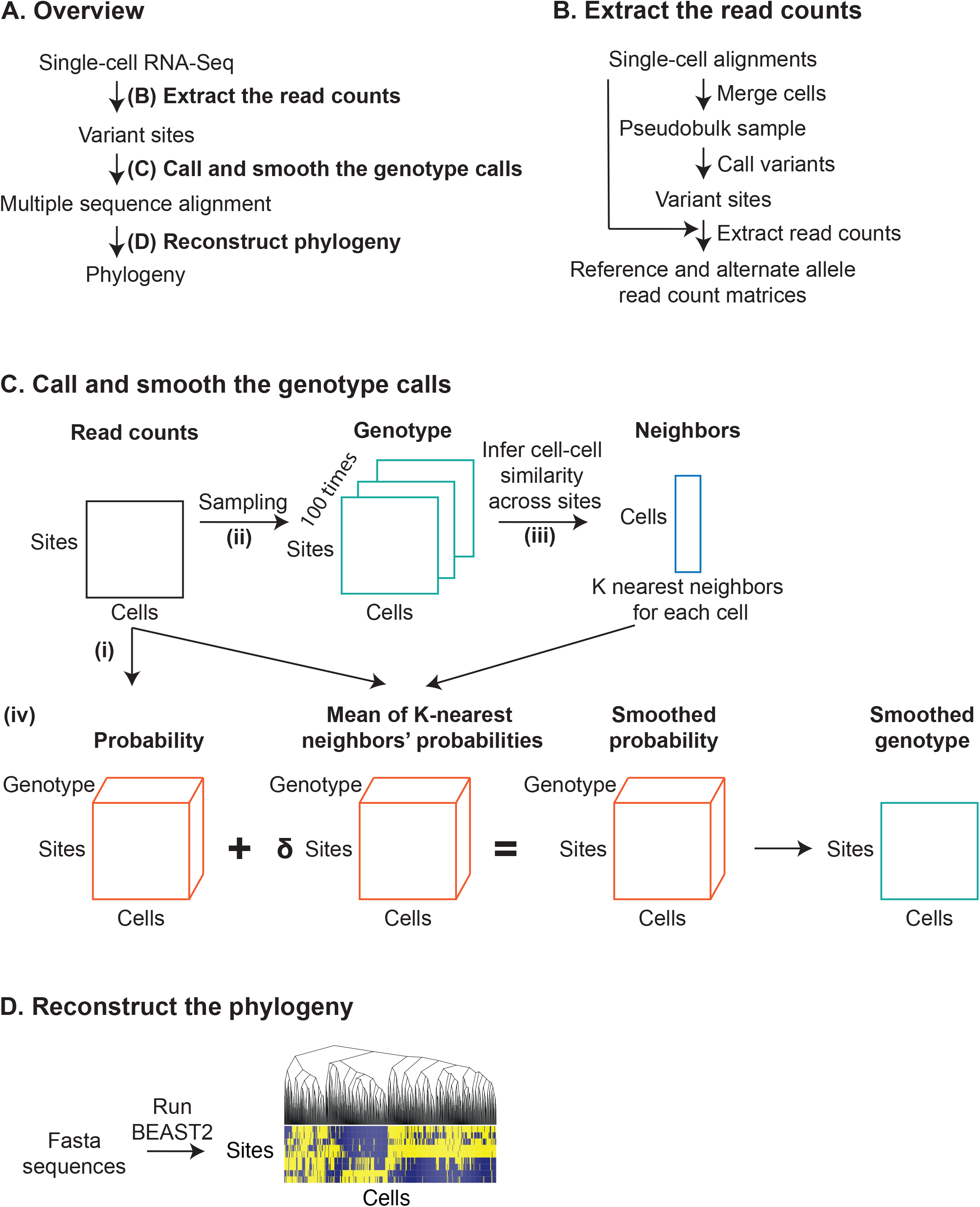
Construction of phylogeny from scRNA-Seq data. (A) Generating a phylogeny from scRNA-Seq alignments consists of three major steps: 1) extracting the read counts, 2) calling and smoothing the genotypes, and 3) reconstructing the phylogeny. (B) We start with the alignments of cells from a single-cell RNA-Seq experiment. To extract matrices of read counts, we identify sites of interest by merging the alignments in a pseudobulk sample and calling variants. Then, at each of the variant sites, in each individual cell, we count the number of reads with the reference and alternate alleles. (C) This shows the genotype calling and smoothing processes. We start from matrices of read counts of reference and alternate alleles seen across sites (rows) in single cells (columns). i) Given the number of reads, we assigned a probability of a (R)eference, (A)lternate, or (H)eterozygous genotype by integrating over a beta-binomial density function. ii) To compare the genotypes of two cells, we sample 100 genotype profiles by drawing from their probability distributions. iii) Comparing every pair of cells leads to a pairwise similarity matrix of genetic distance scores. By looking for the highest scores (excluding itself), we find the *K* nearest neighbors for each cell. iv) With the nearest neighbors, we can smooth the genotype probability of a cell by averaging with the weighted (%) average probabilities of its neighbors. We call the genotype with the highest probability score. (D) We use BEAST2 to infer the phylogeny and produce a final tree using the max clade credibility method.

The algorithm first identified sites that varied across the cells and thus might best reveal phylogenetic structure. To do this, we combined the reads for all the cells into a single pseudobulk sample and then called variant sites using a GATK pipeline (**Figure 1B**). Next, for each cell, we called a genotype (either reference, alternate, or heterozygous) at each of the variant sites (**Figure 1C**). To account for low read depth, genotype calls were smoothed using information from related cells, as described in the Methods. We assigned reference and alternate bases according to the base seen in the alignments, and if the genotype was heterozygous, we assigned an arbitrary surrogate base. Finally, to estimate the phylogeny of the cells, we used BEAST2, a Bayesian phylogenetic inference algorithm (**Figure 1D**).

### Smoothing genotypes enables identification of resistant phenotypes

To verify that genotype smoothing helped recover more accurate phylogenetic relationships, we used a data set where the CAMA-1 breast cancer cell line was experimentally induced to evolve under 6 months of treatment with ribociclib, a CDK4/6 inhibitor, resulting in a resistant cell line (GSE193278) [31]. A sister lineage from the same ancestral population evolved under untreated conditions for the same period and remained ribociclib sensitive. Whole exome sequencing revealed genetic differences between the resistant and parental populations, including nine predicted to be highly deleterious, suggesting that the resistant cells underwent genetic evolution under drug selection [32]. The populations of sensitive and resistant cells could be distinguished by fluorescent labels. Cells were grown for 14 days in both mono- and co-culture, allowing us to distinguish biological differences in the cell populations from technical batch effects.

We pooled data from all cells, processed the data, and obtained 523 variant sites and 400 cells (200 resistant and 200 sensitive). We inferred cancer mutation profiles and phylogenetic trees of the cells’ evolutionary relatedness with or without application of our algorithm for smoothing the genotypes.

Assuming that the cells that evolved under ribociclib selection harbored genetic changes that conferred drug resistance, we expected to observe that resistant and sensitive cells occupied distinct evolutionary branches of the phylogeny. When we compared the mutation profiles generated with and without genotype smoothing, we found that the nucleotide bases called using smoothing better distinguished the sensitive and resistant cells (**Figure 2A**). Without smoothing and imputation, 30% of the values in the matrix were missing, and the genotypic variation showed no association with the resistance status (adjusted Rand Index=0, p=0.44). With smoothing, the genotypes at selected variant sites were significantly associated with the resistance lineage (adjusted Rand Index=0.3, p=0.001), showing that the data captured the evolutionary alterations distinguishing resistant and sensitive cancer lineages.

**Figure 2.**
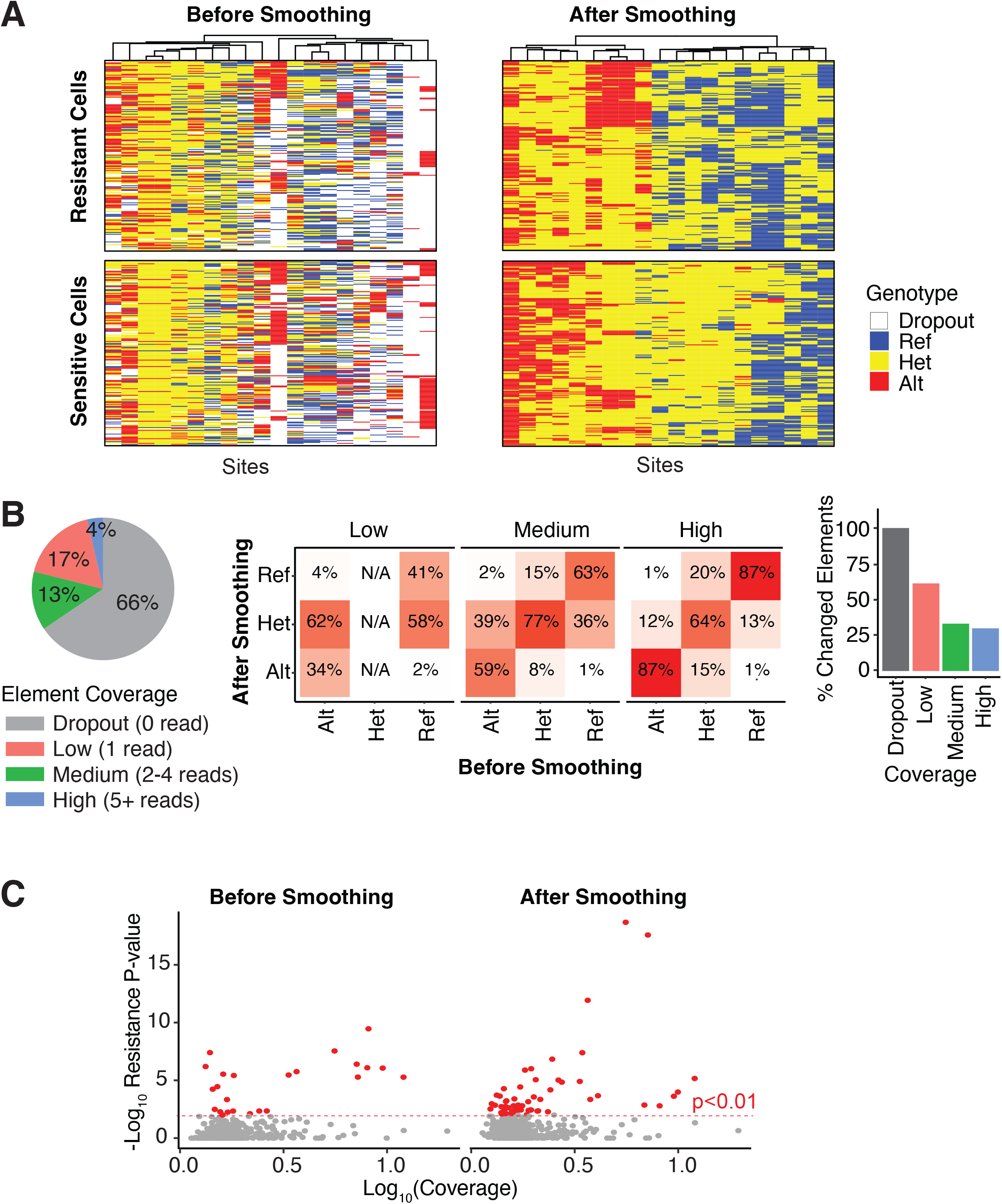
Smoothing the genotype calls. (A) These heatmaps show the genotypes of the resistant (top) and sensitive (bottom) cells for the 20 sites that are most significantly associated with resistance. The heatmaps on the left show the genotypes before smoothing, and the ones on the right show the genotypes after smoothing. (B) (left panel) The pie chart shows the distribution of the number of reads seen in each element in the matrix. (middle panel) The heatmaps show the percent of elements that changed after smoothing. The elements are discretized into low, medium, and high coverage groups. (right panel) The bar plot shows the percent of changed elements in each coverage group. (C) The scatter plots show the association between the mean coverage (x-axis) of each site as well as its association with resistance (y-axis). Sites associated with resistance at p<0.01 are shown in red.

To understand how the smoothing affected the genotype base calls, we examined each element (a specific site and cell in a matrix) of the read count matrices and calculated the percent that were altered by the smoothing (**Figure 2B**). Because smoothing depended on the depth of the coverage (the total number of reads), we discretized the elements into drop-out (0 reads, 66% of elements), low (1 read, 17% of elements), medium (2-4 reads, 13% of elements), and high (>= 5 reads, 4% of elements) coverage. As expected, the genotypes from elements with low coverage were most likely to be altered by smoothing and were changed 63% of the time, in comparison to the ones with medium (34%) and high (30%) reads. The ability to call homozygous genotypes (either reference or alternate) improved with higher coverage. With low coverage, no heterozygous genotypes could be called without smoothing (since only one read was available), and ~60% of the homozygous genotypes were changed to heterozygous after smoothing. For elements with medium coverage, ~40% of the homozygous genotypes were changed; and for high coverage, ~10% were changed.

Finally, to determine if smoothing uncovered more sites associated with the resistance lineage, we examined the association of the 523 genomic sites with resistance status before and after smoothing. After smoothing, more sites showed genetic alterations that were associated with resistance/sensitive lineage status (**Figure 2C**). These results indicated that smoothing enhanced the ability to identify sites linked to the evolutionary divergence of the resistant lineage.

### Phylogenies are not confounded by technical characteristics of scRNA-Seq

A major challenge in analyzing scRNA-Seq data has been the overall low coverage and high drop-out rates [19]. To assess the sensitivity of phylogenetic reconstruction to these characteristics, we performed simulation studies using the CAMA-1 data set.

First, we tested the robustness of the inference to the number of neighbors, *K*, used in the smoothing algorithm (**Figure 3A**). We generated phylogenies after varying *K* from two to 20 neighbors, and scored the association with resistance status using Pagel’s λ, a measure of phylogenetic signal [33]. We found that λ was not dependent on *K*, indicating that the phylogeny was relatively robust to the number of neighbors.

**Figure 3.**
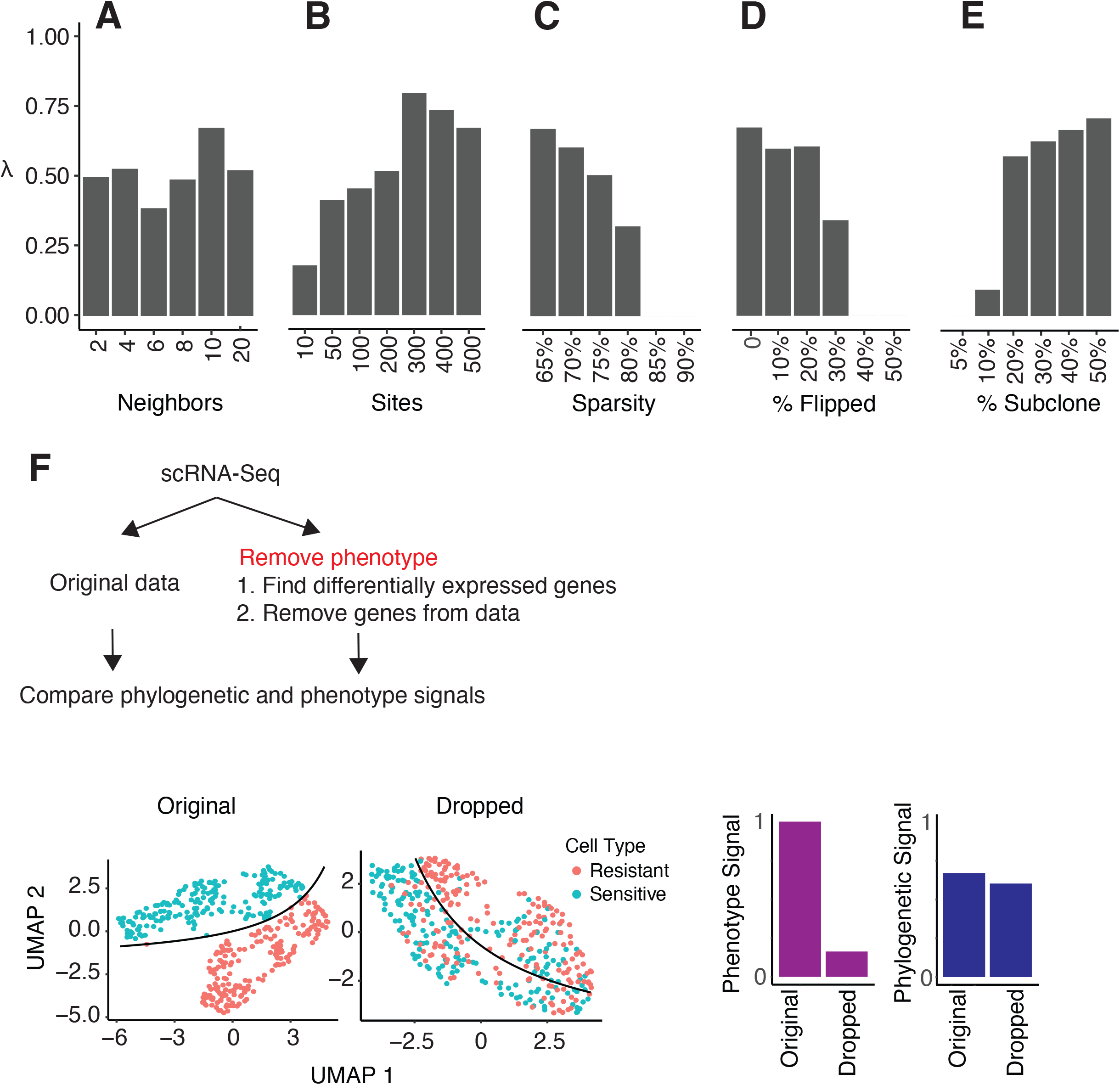
Impact of data quality and parameters on phylogeny. These barplots show the relationship between the phylogenetic signal λ and (A) number of neighbors, (B) number of sites, (C) sparsity, (D) percent of genotype flipped, and (E) subclone size as percent of all cells. (F) The graphic on the top shows the strategy we use to determine whether the phylogenies are confounded by gene expression patterns. The UMAP plots show the resistant and sensitive cells before (left) and after (right) dropping out 1,948 genes associated with resistance. The bar plots represent the resistance signal identified from phenotype (left) and genotype (right) before and after dropping out the resistance-associated genes.

Next, we tested how the number of selected sites impacted the phylogenies (**Figure 3B**). We reconstructed cancer phylogenies after selecting different numbers of genomic sites (10-500) with the highest average coverage. The phylogenetic signal increased as the number of sites increased up to 300 sites, and then began to decrease slowly, potentially due to the addition of noisier data. This suggested, in principle, that more sites should be favored over fewer, although too many low-quality sites can be detrimental. It is known that the number of sites required will increase with the number of evolutionary relationships to be inferred [34]. It is likely that the ideal number of sites depends on the mutational process that is driving the evolution of the cells, and thus may vary from data set to data set.

We also examined how the rate of drop-out affected the phylogenetic signal by taking the unaltered count matrix (with 65% missing values, which we called the *sparsity* of the matrix), and then setting random elements to 0 to increase the sparsity up to 90% (**Figure 3C**). This experiment revealed that the phylogenetic signal decreased with the addition of sparsity, rapidly dropping after 75%. No association of phylogeny with resistance status was seen beyond 85% data sparsity.

Another potential source of noise was the incorrect genotype calls due to random sequencing artifacts or limitations of the imputation algorithm. To test whether these types of errors impacted phylogenetic inference, we simulated additional errors by randomly changing the genotypes of up to 50% of the elements (**Figure 3D**). Up to 20% of the genotypes could be randomized with only a moderate decrease in the phylogenetic signal, but performance dropped drastically with more severe noise.

Next, we tested how well the algorithm could detect low frequency subclones. For this analysis, we constructed data sets consisting of the sensitive CAMA-1 cells, and then spiked in different proportions of resistant cells so that the resistant cell frequencies ranged from 5-50%, while maintaining a constant total cell count (**Figure 3E**). To accommodate small subclones, we modified the filtering criteria and removed sites if over 95%, rather than the default of 90%, of the cells had the same base (see Methods). From analysis of the phylogenies, we found that subclones comprising at least 20% of the cells could be clearly detected, while smaller ones were difficult to detect.

Finally, we tested whether the gene expression patterns affected the phylogenies. Because genotypes could only be assessed (without imputation) from genes that were expressed, it was possible that the expression patterns may have confounded the structure of the phylogenies such that the clades reflected gene expression patterns rather than genotypes. Previously, we reported that resistant CAMA-1 cells had altered gene expression patterns [32] that we confirmed here (**Figure 3F**). We then eliminated the genes with differential expression between the resistant and sensitive lineages (dropping up to 1948 differentially expressed genes with log fold change > 0.05 and detected in >5% of cells) and confirmed that gene expression differences were removed by UMAP. We then generated phylogenies from both the original and scrubbed data sets. After removing the differentially expressed genes, we observed a dramatic decrease in the distinction between the resistant and sensitive cells as quantified by the phenotypic signal (**Figure 3F**). However, the phylogenetic signal remained comparable, showing that the evolutionary divergence of resistant and sensitive cells could be identified from mutational changes, even if there were no differences in the gene expression patterns.

To summarize these analyses, we found that the single cell phylogenies from scRNA-seq data were relatively robust to how our algorithm was applied, but could be affected by the quality of the data. Phylogenies were insensitive to the number of neighbors; more sites (>= 300) and less sparsity (lower is better, with an upper limit of 75%) were better; up to ~20% of poor calls were acceptable; larger subclones were easier to detect, although small ones down to 10% could be found; and the phylogenies indeed reflected the genotype and were not confounded by the gene expression patterns in the data.

### PhylinSic can detect phylogenies across a range of biological conditions

Thus far, the results showed that our algorithm could recover the genetic differences underlying acquired drug resistance in breast cancer cells. To determine whether the method was generalizable and could recover evolutionary relationships produced by other evolutionary forces, we tested it on a range of conditions seen in five public scRNA-Seq data sets where the subclonal structures were determined by independent data sources, including single cell DNA-sequencing or bulk whole genome sequencing (**Table 1**). Each data set contained monophyletic cells that could be grouped into subpopulations with distinct genetics. We reconstructed the phylogenies of cells in each data set and compared them against their previously established subclonal structure. Then, we measured the association between the phylogenies and the subclones using the phylogenetic signal.

**Table 1.** Gold standard data sets.

| <u>Data Set</u> | <u>Description</u> | <u>Sequencing Platform</u> | <u>Evidence of Subclone</u> | <u>Cells</u> | <u>Sites</u> | <u>Sparsity (before filter)</u> | <u>Sparsity (after filter)</u> | <u>Reference</u> |
| --- | --- | --- | --- | --- | --- | --- | --- | --- |
| CAMA1 | ER+/HER2- CAMA-1 breast cancer cells | 10x 3' | Whole exome | 400 | 213,379 | 97% | 65% | bioRxiv doi:10.1101/2021.02.01.429214 |
| N87 | Gastric cancer cell line | 10x 3' | scDNA-Seq | 500 | 12,984 | 92% | 75% | PMID: 32215369 |
| ER+BRCA | Breast tumors, before and after treatment | Fluidigm | Whole genome, scRNA-Seq copy number | 141 | 164,580 | 97% | 53% | PMID: 29093439 |
| MM16 | Multiple myeloma tumors, before and after treatment | Fluidigm | Whole exome, scRNA-Seq copy number | 46 | 144,057 | 94% | 32% | PMID: 31558476 |
| MM34 | Multiple myeloma, primary and metastasis | Fluidigm | Whole exome, scRNA-Seq copy number | 127 | 326,089 | 95% | 16% | PMID: 31558476 |

We begin by investigating more deeply the CAMA-1 breast cancer cells from above [32]. In this data set, three samples were sequenced: one sample with resistant cells grown in monoculture, one with sensitive ones in monoculture, and one where the resistant and sensitive cells were co-cultured. After combining the cells from these three samples, we randomly selected 200 sensitive and 200 resistant cells and reconstructed their phylogeny. The phylogeny bifurcated into two major clades, where one was predominantly composed of resistant cells, and the other was mainly sensitive (phylogenetic signal *λ* =0.67, p=2.8e^-23^) (**Figure 4A**). There was no difference in the distribution of the monocultured or co-cultured cells (phylogenetic signal *λ* =0.28, p=0.001), indicating that the phylogeny reflected genetic heterogeneity and was not confounded by environmental or batch sampling effects.

**Figure 4.**
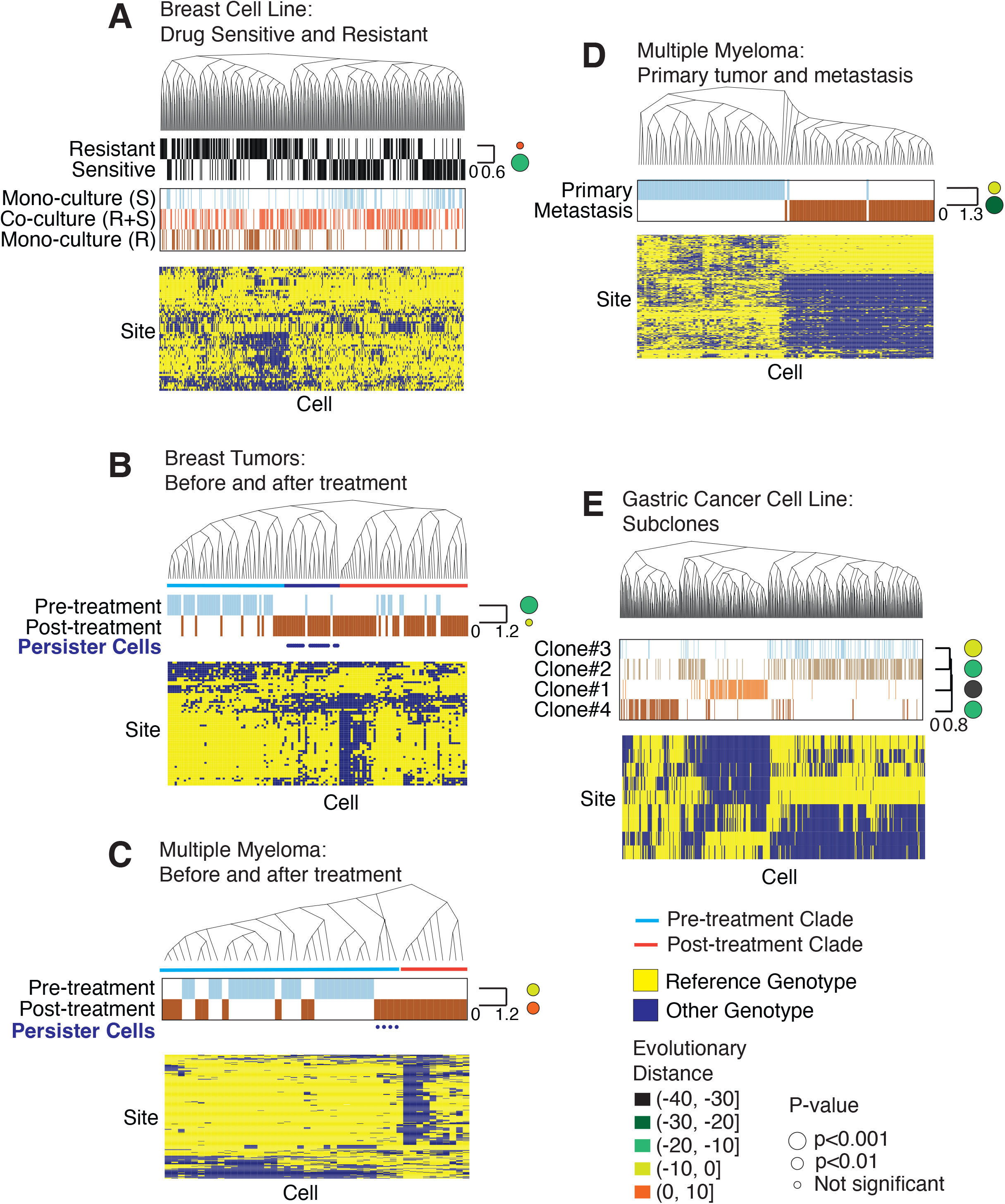
Associations of phylogenies with previously identified subclones. We have generated phylogenies from five scRNA-Seq data sets. The evolutionary distance for each condition is shown as the circles at the right of the heatmap. The color and size of the circles represent the evolutionary distance and statistical significance. (A) These cells were derived from the CAMA-1 cell line grown in three separate cultures, ribociclib sensitive (S) and resistant (R) cells cultured by themselves, or mixed and cultured together (R+S). The cells induced to be drug resistant are shown as black lines in the *Resistance* bar. The genotypes of the sites with high association with the phylogeny are shown in the heatmap at the bottom. (B) This data set consists of tumor cells collected from ER+ breast cancer tumors before and after treatment. The color bars show the time point of each cell. The phylogeny is divided into clades of pre- (blue) and post-treatment (red) genotypes. Persister cells are delineated with dots. (C) This contains tumor cells from a multiple myeloma patient before and after chemotherapy treatment. The phylogeny is divided into clades of pre- (blue) and post-treatment (red) genotypes. Persister cells are indicated with dots. (D) This data set contains cancer cells collected from the primary or metastasis site from a multiple myeloma patient. (E) This shows cells from the N87 gastric cancer cell line. Four subclones were previously reported.

We then tested the ability of PhylinSic to infer genetic relationships from patient tumor data. In one patient derived tumor data set, we studied metastatic ER+ breast cancer tumors [35] with cells profiled both before and after treatment with chemotherapy (**Figure 4B**). From this data set, we studied the patient evolutionary relationships of cancer cells in the tumor with the clearest evidence of subclone architecture (patient 3). In another data set, we analyzed tumors from patients with multiple myeloma [36] (**Figure 4C**). In both cases, the distribution of the pre- and post-treatment cells were associated with the phylogeny (breast *λ* = 0.73, p=1.5e^-17^, melanoma *λ* = 1.1, p=1.1e^-6^), indicating a shift in the genetics of the tumors after treatment. For the breast cancer tumors, a minority of the pre-treatment cells (23%) shared a genetic lineage with the post-treatment cells, indicating that some pre-treatment cells had a pre-existing resistant genotype, as we previously reported [35]. This pattern was not observed in the multiple myeloma tumors, although it was possible that too few cells were sequenced to detect this population.

To further characterize the evolutionary distinctness of pre- and post-treatment cells, we calculated the evolutionary diversity of cells from each population across the phylogeny. We then assessed whether cells of each population were phylogenetically clustered within specific branches of the evolutionary tree, using standard-effect-size mean pairwise distance (SES MPD). Negative evolutionary distances indicated phylogenetic clustering while positive distances indicated phylogenetic evenness. In both breast cancer and myeloma data sets, post-treatment cells were more evenly distributed across the phylogeny than pre-treatment cells, indicating that post-treatment cells were more genetically diverse despite the effects of drug selection.

In both breast and multiple myeloma tumors, we observed distinct pre- and post-treatment clades (**Figure 4B, 4C**). Interestingly, in both tumors, a small set of cells that survived after treatment genetically resembled the pre-treatment genotype (inferred to be drug sensitive). That is, these *persister* cells were closely related genetically to the sensitive pre-treatment cells, and were more distantly related to the resistant post-treatment cells (**Figure 4B, 4C**). This suggested that the local tumor microenvironment provided resistance to these otherwise sensitive cells (e.g., by limiting drug penetration). Additionally, pre-treatment we observed a small population of cells in the breast tumor that were genetically similar to the dominant post-treatment lineage (resistant cells), indicating the existence of resistant cancer genotypes within the heterogeneous tumor prior to treatment (**Figure 4B)**.

In addition to evolution driven by drug treatment, we examined the evolution of primary and metastatic tumors from a multiple myeloma patient [36] and found that the cells from the two tumors were nearly completely separated into distinct lineages (**Figure 4D**). A small number of cells within the primary tumor belonged to the lineage that dominated in the metastasis, but the dominant lineage of the primary tumor was completely absent of metastatic cells. This result suggested that a single (or small number of related) genetic lineage(s) colonized the distant location and produced a distinct genetic population at the distant site.

Since PhylinSic could identify genetic changes resulting from drug treatment and metastasis, both of which were strong selective events, we finally tested whether it could detect differences resulting from more subtle genetic drift. To do this, we applied it to scRNA-Seq profiles of the N87 gastric cancer cell line that was cultured under conditions without any specific selective pressure [37]. We chose this cell line because, in the published study, it was the most deeply characterized. There, four subclones were identified by scDNA-Seq and scRNA-Seq. In the PhylinSic-derived phylogeny, distinct clades could be seen for Clone 1 (*λ*=1.1, p=8.8e^-28^) and Clone 4 (*λ*=0.68, p=1.8e^-79^), while we found some disagreement in the classification of Clones 2 (*λ*=0.5, p=5.0e^-30^) and 3 (*λ*=0.34, p=1.8e^-15^) which were more difficult to distinguish (**Figure 4E**). The mutations that we identified supported the PhylinSic classification although some mutations detectable by scDNA-Seq may not have been observable in the scRNA-Seq.

Taken together, these results demonstrated that PhylinSic could reconstruct the phylogenetic relationships, enabling biological insight of cells across a range of biological settings (*in vitro* and *in vivo* throughout treatment or across metastatic sites) and sequencing platforms.

### Integrating genetic and phenotypic inference

Having seen the ability of PhylinSic to reconstruct the genetic relationships across single cells from scRNA-Seq data, we next sought to link these genotypes with the phenotypes of the cells, as represented by gene expression profiles. To do this, we analyzed the breast tumor data set (represented in Figure 4B) to determine how phenotypes evolved in response to drug treatment.

To determine the phenotypic traits under evolutionary selection, we measured the phylogenetic signal λ for the Hallmarks pathways [24]. In the breast cancer data set, this revealed the phenotypes that were significantly dysregulated in each of the 5 major cancer clades (clades A-E). This included pathways previously linked with breast cancer including Wnt/β-Catenin signaling (false discovery rate, fdr=0.005), K-Ras (fdr=0.001), estrogen response (fdr=0.035), and mitotic spindle (fdr=0.002) (**Figure 5A, 5B**). Many of these pathways were upregulated in only one clade of the phylogeny, predicting the event that led to activation of the pathway. For instance, one clade composed almost entirely of pre-treatment cells (clade A in Figure 5A) had a high estrogen response signature, which was low in the remaining clades of post-treatment cells (clades B-E). Since these tumors were ER+, this predicted that there was loss of estrogen dependency after treatment, which was driven by a genetic change. In contrast, β-catenin signaling, a pathway correlated with drug resistance in a number of contexts [38, 39], was high in the post-treatment cells.

**Figure 5.**
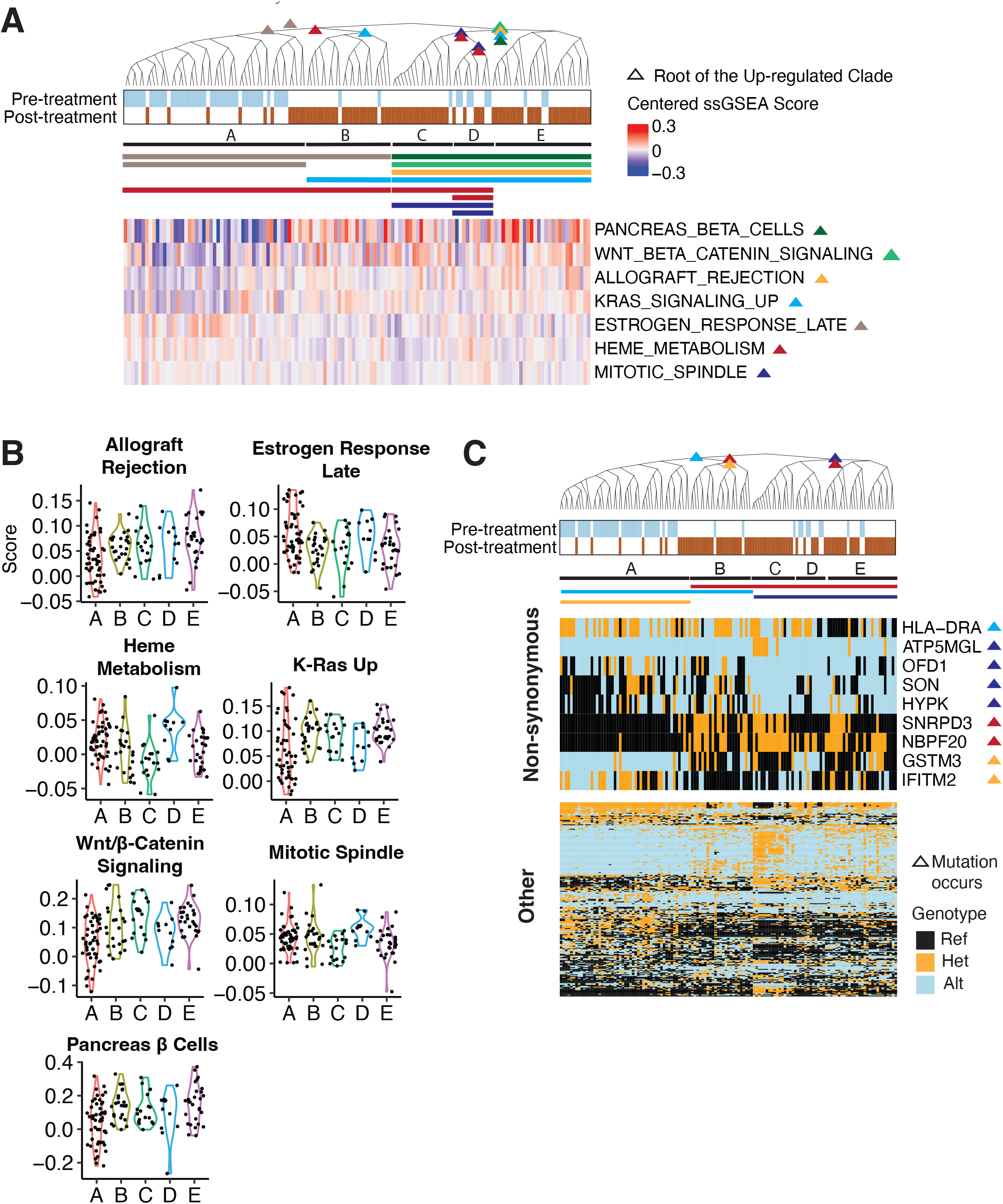
Associating the genotypes with phenotypes. (A) The phylogeny from the breast tumor data set in figure 4B is reproduced here with new annotations. The phylogeny is split into five clades (A–E). The clades associated with high pathway scores are marked with a colored triangle. The siblings of these clades have low pathway scores. The heatmap on the bottom shows the mean-centered ssGSEA scores for the Hallmark pathways significantly associated with the phylogeny based on phylogenetic signal (fdr<0.05). Each pathway is labeled with colored triangle(s) that indicate the clade(s) with high pathway score. The phylogenetic associated clades are marked by the horizontal bars below the phylogeny. (B) The violin plots show the ssGSEA scores for the five clades. (C) This phylogeny and clades are annotated similarly to those in figure A, except that gene mutations are associated with the phylogeny, rather than pathway scores. The heatmap on top shows genes with non-synonymous mutations that are correlated with the phylogeny. The bottom heatmap shows the remaining genes that are correlated with the phylogeny (fdr<0.01).

Finally, in this data set, we noticed a set of persister cells, noted above, with a sensitive genotype, that were retained after treatment (clade B in Figure 5A). While these cells were genotypically distinct from the other post-treatment cells, we find here that some aspects of their phenotypes were similar (**Figure 5B**). Notably, they shared high K-Ras and β-catenin pathway scores with the other resistant cells. However, while the other resistant cells had a genetic association with β-catenin, the persisters had a genetic association with K-Ras. While the post-treatment cells had high activities for both pathways, the genetics underlying those pathways were different.

To determine whether any recognizable mutations were associated with these phenotypes, we examined the 1000 mutation sites that were used to construct the phylogeny and measured their phylogenetic signal at each internal node of the phylogenetic tree (**Figure 5C**). We found 228 sites that were significantly associated with the phylogeny (fdr<0.01). However, the vast majority occurred in non-coding regions and thus were not likely to be driver mutations. We focused on the nine non-synonymous mutations and found one, GSTM3 (NM_000849, p.V224I), which had a significantly different genotype in the post-treatment cells. GSTM3 was a β-catenin target that had been linked to chemoresistance and other phenotypes in cancer [40-42]. While the phylogeny did not determine whether this was a driver mutation activating a β-catenin signature, it nevertheless provided a novel model of the biology and revealed hypotheses to be tested.

## Discussion

We have developed a method, PhylinSic, to reconstruct evolutionary relationships within tumors at the single-cell resolution and to link genotype to phenotype using scRNA-Seq data. We demonstrated the potential that this had for generating biological insight. Using probabilistic approaches, we addressed the challenges inherent in the platform, such as low read coverage, drop-out, and biases in the gene expression patterns. The approach we developed to call bases could also be applied to other contexts requiring accurate base calls from scRNA-Seq, such as estimates of mutation burden or calling of tumor vs. normal cell types. Finally, we had shown that our method, PhylinSic, could generate plausible phylogenetic trees on a range of data sets, from cell lines to patient tumors, and across a range of scRNA sequencing platforms including 10X Genomics and Fluidigm.

For drug resistance (in a CDK4/6 inhibitor and chemotherapy setting), we saw evidence for pre-existing populations of resistant genotypes, even pre-treatment. Intriguingly, in both tumor drug treatment data sets, we saw two distinct lineages of cells that survived after treatment. This result suggested that resistance mechanisms may be commonly multiclonal. Further, the phenotypic data showed that the post-treatment breast tumor cells, regardless of the lineage, had activation in at least two signaling pathways: K-Ras and β-catenin. While all resistant lineages had increased activation of those pathways, the differing genetic backgrounds suggested that they acquired activation through different mechanisms. This possibility implied that inhibition of multiple distinct mechanisms may be necessary and thus supported the hypothesis that heterogeneity is a mechanism of resistance, and multiple (or all) mechanisms would need to be targeted to achieve a durable response Beyond subclones, a full phylogenetic model provided the genetic relatedness of each cell, quantified from evolutionary distances that reflected the underlying mutation rates and divergence times. This allowed principled (evolutionary) associations between genotype and phenotype that enabled us to distinguish the traits associated with evolution, and also determine whether those traits were evolved multiple times independently through convergent evolution–a situation that may confound cancer treatment [43].

However, there remain limitations in our approach that need to be addressed. Phylogenetic models, in principle, can reveal the timings of historical evolutionary events that could be used related to events in the life of the patient (e.g., drug selection, time of metastasis, immune surveillance). However, this requires a carefully calibrated clock model that can contend with possible differences in the evolutionary rate across cancer cell lineages, e.g., with differing DNA damage repair abilities. This could be addressable with a model that integrates the mutational processes that drive the rate of mutation in each lineage [44]. This would also reveal how differing rates or processes of evolution along different lineages can impact the malignancy of the cells. In any case, the lens of evolution reveals not only the life history of the tumors, but provides an understanding of the current genetic state of the tumors, and potentially vulnerabilities in their evolutionary trajectory.

## Acknowledgments

This research was supported by the National Cancer Institute of the National Institutes of Health (NIH) under awards number U54CA209978 (AHB, JTC) and U01CA264620 (AHB), as well as NIH award UL1TR003167 (JTC).

